# Protein shape sampled by ion mobility mass spectrometry consistently improves protein structure prediction

**DOI:** 10.1101/2021.05.27.445812

**Authors:** SM Bargeen Alam Turzo, Justin T. Seffernick, Amber D. Rolland, Micah T. Donor, Sten Heinze, James S. Prell, Vicki Wysocki, Steffen Lindert

**Affiliations:** Department of Chemistry and Biochemistry and Resource for Native Mass Spectrometry Guided Structural Biology, Ohio State University, Columbus, OH, 43210; Department of Chemistry and Biochemistry and Materials Science Institute, University of Oregon, Eugene, OR, 97403

## Abstract

Among a wide variety of mass spectrometry (MS) methodologies available for structural characterizations of proteins, ion mobility (IM) provides structural information about protein shape and size in the form of an orientationally averaged collision cross-section (CCS). While IM data have been predominantly employed for the structural assessment of protein complexes, CCS data from IM experiments have not yet been used to predict tertiary structure from sequence. Here, we are showing that IM data can significantly improve protein structure determination using the modeling suite Rosetta. The Rosetta Projection Approximation using Rough Circular Shapes (PARCS) algorithm was developed that allows for fast and accurate prediction of CCS from structure. Following successful rigorous testing for accuracy, speed, and convergence of PARCS, an integrative modelling approach was developed in Rosetta to use CCS data from IM experiments. Using this method, we predicted protein structures from sequence for a benchmark set of 23 proteins. When using IM data, the predicted structure improved or remained unchanged for all 23 proteins, compared to the predicted models in the absence of CCS data. For 15/23 proteins, the RMSD (root-mean-square deviation) of the predicted model was less than 5.50 Å, compared to only 10/23 without IM data. We also developed a confidence metric that successfully identified near-native models in the absence of a native structure. These results demonstrate the ability of IM data in *de novo* structure determination.

## Introduction

Proteins are at the core of virtually all cellular processes. Therefore, comprehensive knowledge of protein structures with atomistic detail can be beneficial for several pharmaceutical applications such as vaccine design^1^, drug discovery^2, 3^, enzyme design^4^, self-assembling molecular machines^5^, and many more^6^. Mass spectrometry (MS) has become a prominent technique in the field of structural biology due to its ability to provide structural information for proteins and protein complexes. MS can be particularly beneficial because it is faster, can work for heterogeneous samples, can be used routinely at all stages of a project, and has fewer sample preparation complications compared to high-resolution techniques such as X-ray crystallography, cryo-electron microscopy (cryo-EM) and nuclear magnetic resonance (NMR) spectroscopy. Several findings for protein structures in the gas phase also suggest that features such as elements of secondary structure, compactness and quaternary structure can be preserved during the transition from solution to desolvated state^7–9^. For these reasons, structural MS can be very beneficial particularly when high-resolution methods are not feasible^10, 11^. Various methods have been developed and coupled to MS to study protein structures^12, 13^ such as chemical crosslinking (XL)^14^, covalent labeling (CL)^15^, surface induced dissociation (SID)^16^ and other ion activation methods such as collision-induced dissociation (CID), electron capture/transfer dissociation (ExD), and ultraviolet photodissociation (UVPD), hydrogen-deuterium exchange (HDX)^17^ and ion mobility (IM)^18^. While such MS techniques may provide diverse details and routine analysis of structures, experimental data collected from experiments are sparse and cannot unambiguously determine atomic-resolution structure^19^.

An alternative approach to experimental structure determination is to use computational modelling methods. These approaches, such as structure prediction from sequence or protein-protein docking, can also provide insight into atomistic details of biomolecules but are frequently limited in accuracy due to the large conformational sampling space among other challenges^20^. While these methods can be successfully utilized in the absence of experimental data, sparse experimental data are often used to guide and improve modeling^19, 21, 22^. Experimental data from various MS techniques have recently proved pivotal within integrative structural biology frameworks ^14, 17, 23–40^.

In IM, ions are transferred into an inert gas chamber at a constant pressure and temperature under the influence of a weak electric field^41, 42^. This technique is regularly utilized to separate protein structures based on their shape and size. IM can also provide a rotationally averaged collision cross section (CCS_IM_) of the protein which is related to the amount of momentum exchanged between ion and buffer gas over the course of the collisions and can be thought of as somewhat like rotationally averaged cross sectional area^43^. Several methods have been developed to predict CCS from protein structure. Among these, the most physically realistic algorithms are the trajectory method (TJM)^44, 45^ and diffuse trajectory method (DTM)^46^ which integrate Newton’s equation of motion to calculate the classical scattering of gas particles. Both TJM and DTM explicitly account for long-range interactions through Lennard-Jones potentials to approximate momentum transfer from each gas particle to the collided ion. CCS obtained from these methods is extremely accurate^45^, but these calculations can be slow. Due to the high computational cost, prediction methods such as elastic hard sphere scattering^47^, projection superposition approximation (PSA)^48^, local collision probability approximation^49^ and projection approximation (PA)^43^ make further approximations on TJM, resulting in faster CCS calculations. Among these approximated methods, PA is the simplest and fastest, because it neglects the scattering and long-range interactions^43, 50^. CCS_PA_ only accounts for the collisions of a gas particle with the ion based on hard sphere atomic radii by calculating the average cross-sectional area of the protein structure as experienced by the buffer gas. Using the CCS projection approximation calculation tool IMPACT, calculations are about 10^6^ times faster^43^ than the most rigorous methods and are widely used for comparison with experimental IM data. Therefore, PA approaches are advantageous for use in integrative modeling, where the CCS calculation is required for thousands of structures that are obtained from Monte Carlo sampling.

Several instances of structural modeling in conjunction with IM data have been reported. IM spectra have been successfully predicted with the structure relaxation approximation (SRA) method^9^. This method uses molecular dynamics simulations to model structures in the specific charge states. It then utilizes CCS_PSA_ of the generated structures to predict an overall IM spectrum. The SRA method indicated that systems studied with IM methods are generally consistent with retention of many residue-residue contacts determined by X-ray crystallography. Furthermore, IM data have been incorporated in computational modelling for protein complex structure prediction. In these methods, coarse-grained models generated using the Integrative Modelling Platform^51^ were ranked and clustered based on the agreement between predicted and experimental CCS measured from IM^28, 35^. CCS_IM_ values for complexes and their individual subunits have also been successfully used to approximate rough intersubunit distance used as restraints in modeling methods to identify coarse-grained topologies of complexes^36–39^. In addition to complex structure prediction, work has also been done to show correlation between IM data and structural similarity (RMSD)^9^. While several studies have demonstrated that IM data can be predicted and utilized with various computational methods, IM data have not yet been utilized to predict tertiary structure from sequence.

Therefore, in this work a non-stochastic grid-based algorithm, PARCS, has been implemented in Rosetta^52, 53^ to predict CCS from structure. It has been demonstrated that PARCS yields comparable results to IMPACT in terms of speed and accuracy. Next an IM score term has been developed for use in the *ab initio*^54–56^ and comparative modelling (CM)^57^ protocols in Rosetta, in combination with the Rosetta all-atom scoring function^58^. This score term scored structures based on their (dis)agreement with experimental IM data. When this score term was included, the prediction of structures improved for a benchmark of 23 proteins: the RMSD improved by an average of 2.01 Å and 15/23 structures were predicted accurately.

## Methods

### Projection approximation using rough circular shapes

Average CCS of biomolecules are determined from IM experiments based on the amount of time required for the ion to traverse the region of inert buffer gas (usually helium or nitrogen) under the influence of a weak electric field^43, 45^. To use IM data in a structure prediction protocol, we developed Projection Approximation using Rough Circular Shapes (PARCS) in Rosetta. The schematic (A) and the illustration (B) in Figure 1 demonstrate how the PARCS algorithm computes CCS from structure and estimates area of a projection, respectively. The PARCS algorithm, as shown in Figure 1A, takes 3D atomic protein coordinates as input. Next, the structure is randomly rotated. For each rotation, the structure is projected on a 2D grid (grid cell area of 1 Å^2^) in the *x-y*, *x-z*, and *y-z* planes as shown in Figure 1A (i). In the 2D grid, the projection of the protein is centered, and the grid extends 5 Å beyond the most extreme atom in each direction. For each atom on the 2D grid (Figure 1A(ii)), the center grid cell is filled as denoted by the blue grid cell in Figure 1B (ii). Then, eight additional cells (red grid cells in Figure 1B (ii)) are also filled. The distance of these eight grid cells from the central cell (i.e., radius of the circular projection) is based on the sum of the radii of the projected atom and the buffer gas (*r* in Figure 1B (ii)). An effective atomic cross-sectional radius of 1.91Å is used for heavy atoms (carbon, sulfur, oxygen, nitrogen and phosphorous) and 1.21 Å is used for hydrogen atoms. A buffer gas radius of 1.0 Å and 1.82 Å is used in the case of helium^43^ and nitrogen^59^, respectively. The eight points are positioned such that two adjacent points on the circumference form a 45° angle from the center point as shown in Figure 1B (ii). This process is repeated for all atoms in the protein, filling the overall grid as shown in Figure 1B (iii). Finally, the projection area (*A*) is derived by summing the areas of the filled grid cells. From the *x-y*, *y-z*, and *x-z* projections for each random rotation *i*, three projection areas 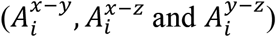 are obtained. The CCS of the structure (*CCS_PARCS_*) is then acquired from the average area of the total number of projections (N=3R, where *R* is the total number of random rotations) as shown in Equation 1.

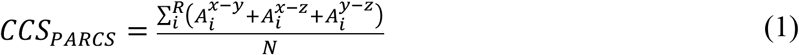

**Figure 1:**
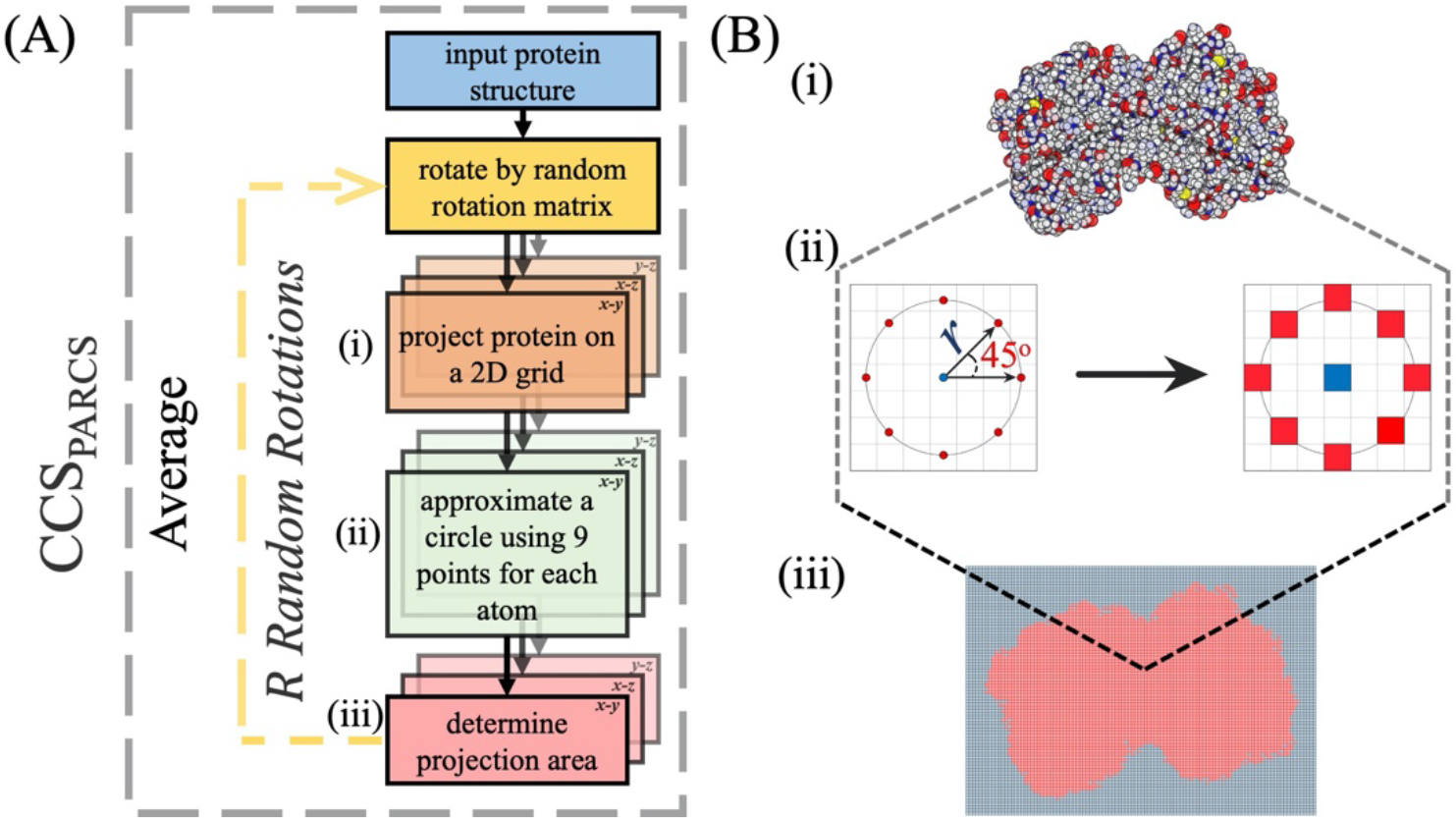
(A) Schematic of the PARCS algorithm to predict CCS from structure. (i) Three projections are obtained from each rotation. (ii) For each atom in each projection the 2D grid is filled according to a 9-point circle approximation. (iii) The projection area is determined from the number of filled grid cells. (B) (i) Illustration of a 2D projection of a single random rotation where the carbon, sulfur, oxygen, nitrogen, and hydrogen are colored grey, yellow, red, blue, and white respectively. (ii) Each atom is projected on a grid with a cell size of 1Å2. The center grid cell and eight other grid cells at a distance *r* (based on the radii of the given atom and the buffer gas) from the center of the atom are filled. (iii) Projection of the randomly rotated protein after the grid cells are filled according to the PARCS algorithm.

### IM score function in Rosetta

CCS from experimental IM data were incorporated as a spatial restraint for integrative Rosetta modelling as it provides information about protein size and shape. Therefore, to integrate this information in Rosetta for protein structure prediction, a score term (*IM_Score_*) was developed to quantify agreement of protein structures with IM data, using CCS as the restraint. The evaluation score, *E_IM_*, was defined as a sum of the *IM_Score_* score term with the Rosetta REF2015 score function^58^ as shown in Equation 2.

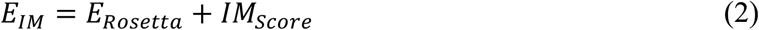

In Equation 2, *E_Rosetta_* is the energy of the structure obtained from the Rosetta REF2015 score function. The IM_Score_ term is a penalty function (as defined and shown in Equation 3 and Figure S1, respectively) based on the absolute difference (ΔCCS) between CCS_PARCS_ and CCS_IM_. This function includes a lower bound (LB) and an upper bound (UB) cutoff (as shown in Equation 3) to account for error^24^. ΔCCS below LB (10Å^2^) are not penalized and ΔCCS above UB (100Å^2^) are given a maximum penalty of 100, with a fade function used in between. Conceptually, this scoring function penalizes structures with high deviation from experiment.

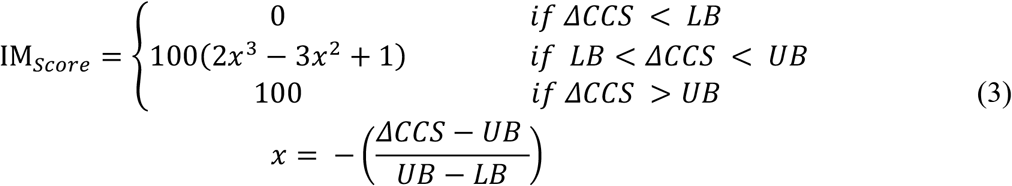

### IM datasets

In this work, our aim was to study predominantly globular and ordered proteins within all datasets. Values from CCS_PARCS_ were compared to CCS_IMPACT_ as well as evaluated for speed and precision on 4465 non-homologous protein structures (PARCS evaluation dataset) extracted from the protein databank (PDB)^60^ (http://www.rcsb.org/) using the PISCES^61^ webserver (http://dunbrack.fccc.edu/pisces). For this dataset, the sequence identity was less than or equal to 10%, sequence length was between 40 – 250 residues, non-X-ray and CA-only entries were excluded and the PDBs were culled by chain. For CCS prediction and speed comparison, PARCS was benchmarked against IMPACT^43^ (with flag ‘-H’ to include hydrogens) based on the calculations performed on the PARCS evaluation dataset. This dataset was also used to test the convergence of PARCS with respect to the number of rotations. In this convergence test, the standard deviation of 100 separate CCS calculations for each protein at varying numbers of rotations were obtained and assessed for the optimal number of random rotations required for calculations to converge.

To evaluate the ability of the score term (Equation 3) to distinguish native from non-native protein models in the case of an error-free CCS prediction, a set of 100 proteins was randomly selected from the PDB (list of monomers shown in Table S1) using the PISCES webserver, where the sequence length ranged from 24 to 154. A set of structure prediction experiments (which will be described in detail in the following sections) was performed on this dataset, where the experimental CCS was simulated by predicting CCS of the native structure with PARCS. Therefore, this dataset was referred to as the ideal dataset. The simulated CCS values ranged from 595 Å^2^ to 1710 Å^2^ for the 100 proteins in the ideal dataset. The score function was also tested on actual experimental IM data, i.e., structures with CCS_IM_ (experimental dataset). The experimental dataset^18, 62–66^ consisted of 23 monomeric proteins that also had structural information deposited in the PDB (as outlined in Table S2). Sequence lengths ranged from 26-691 residues and CCS_IM_ values (for the lowest charge states) ranged from 588 Å^2^ to 4580 Å^2^. Additionally, these proteins exhibited an average percent disorder of only 13% as calculated by the Rosetta ResidueDisorder^67, 68^ application.

### *Ab initio* and comparative modeling protocol for structure prediction

To test whether shape and size information encoded in IM data were sufficient to discriminate between low and high RMSD models of single-subunit proteins, we tested our algorithm on both the ideal and experimental dataset. For these two datasets, the Rosetta *ab initio* protocol was used for proteins with sequence length less than 155 residues, otherwise the Rosetta multi-template comparative modeling (CM) protocol was used. The templates and weights associated with all proteins for CM are provided in Table S4. The 3mers and 9mers fragments required for both protocols were generated using the fragment picker tool^69^ in Rosetta. The protocols (*ab initio* and CM) for both the ideal and experimental data set are further detailed in the SI. All structures generated from the *ab initio* and comparative modeling protocols were subjected to the Rosetta Relax protocol. The IM data, ideal and experimental, were then used to score all the structures generated for each protein in Table S1 and Table S2, respectively. The top scoring model was designated as the predicted structure.

### Analysis metrics used for evaluating predictions

We quantitatively assessed the quality of our predicted models using several of the following metrics. The global RMSDs (root-mean-square deviations) of the predicted models to their native structures were calculated. Predictions with IM data where RMSD was within 0.5 Å of the RMSD of the structure predicted without IM data were defined as unchanged. Next, P_near_ ^70^ a goodness-of-energy funnel metric (at *k_B_T* and *λ* set to 4 and 3 Å respectively), was used to compare the score versus RMSD distributions predicted with and without IM data. P_near_ ranges from 0 (a poor energy funnel) to 1 (a well-defined energy funnel). Predicted structures from both ideal and experimental datasets were evaluated with these two metrics. The predicted structures from the experimental dataset were further evaluated with the template modeling score (TM-score)^71^ and global distance test total score (GDT_TS)^72^. Template modeling score (TM-score) was used to assess the topological similarity of the predicted structures to native structures using the TM-score program^71^. The TM-score metric ranges from 0 to 1, where scores below 0.17 indicate randomly chosen unrelated proteins and a score higher than 0.5 corresponds to structures being generally in the same fold and a score of 1 indicates a perfect match. Global distance test total score (GDT_TS) values were used to assess the optimal superposition between the predicted and the native structure by identifying groups of residues in the predicted model that differ from that of the native structure by less than a distance cutoff. GDT_TS for the Cα atoms at a distance cutoff of 5 Å was calculated using the LGA program^72^. GDT_TS values range from 0 to 100 where higher GDT_TS is indicative of a predicted structure being similar to the native structure.

### Confidence metric used for identifying accurate and inaccurate predictions

A metric was developed to quantify confidence in predictions in the absence of known structure. This confidence metric was defined as the average per-residue score of the top 100 scoring models predicted with IM data (using Equation 2). The specific metric was chosen because lower scores per residue are generally associated with more native-like structures. Thus, structures were defined as high confidence if the average residue score was less than −2.54 (above which structures were defined as low confidence). Instances where the RMSD of the prediction was less 5.50 Å and average residue score less than −2.54 were considered successful confidence measure cases. We chose an RMSD cutoff of 5.5 Å since below that RMSD, protein topologies are generally predicted correctly.

## Results and discussion

In this study, to utilize IM data to predict tertiary (monomeric) structures in Rosetta, an algorithm designed for rapid prediction of CCS has been developed and implemented. This method uses Projection Approximation via a grid-based calculation of Rough Circular Shapes (PARCS). Subsequently, a score function was developed that assessed the agreement of Rosetta-generated models to the CCS_IM_ for tertiary structure prediction.

### Calculation of CCS using PARCS was fast, accurate, and results were comparable to existing software

Area calculation in projection approximation methods is typically performed using Monte Carlo integration methods. In such an approach, probes representing the buffer gas particle are fired upon the randomly oriented 2D-projected target structure to calculate the area of the projection. A large number of probes is usually required for CCS calculations to converge. However, when a large number of probes is used, random probes frequently survey areas with no protein present, resulting in unnecessary calculations and thus adding to the computational cost^73^. Therefore run-to-run variability in probe-based projection area calculation per rotation is common. To circumvent this issue, in PARCS, the projection area is calculated by projecting the structure on a 2D grid and then geometrically estimating the projection area directly (by geometrically filling the grid based on locations of atoms and radii of atoms and probes). This approach eliminates the variability in projection area calculation. Therefore, the only attribute contributing to the variability in CCS calculations using PARCS is the random rotations.

To benchmark our PARCS algorithm, CCS values for 4465 non-homologous protein structures in the PARCS evaluation dataset were calculated. Results for convergence of CCS calculations at varying number of random rotations on the PARCS evaluation dataset are shown in Figure 2A. The average standard deviation of the CCS distributions for 100 rotations was only 2.26 Å^2^ (which was less than 0.2% of the CCS_PARCS_ on average) and decreased as the number of rotations increased. The average of the standard deviations of the CCS distributions was well below 2.0 Å^2^ for more than 100 rotations as shown in Figure 2A and summarized in Table 1. For CCS_PARCS_, the default number of rotations was set to 300, where the average standard deviation of the distribution was 1.31 Å^2^.

**Table 1:** Average standard deviation of CCS calculations for proteins in the PARCS evaluation dataset at various rotations.

| Rotations | Avg std dev (Å<sup>2</sup>) |
| --- | --- |
| 100 | 2.26 |
| 150 | 1.85 |
| 200 | 1.60 |
| 250 | 1.42 |
| 300 | 1.31 |
| 350 | 1.22 |
| 400 | 1.16 |

**Figure 2:**
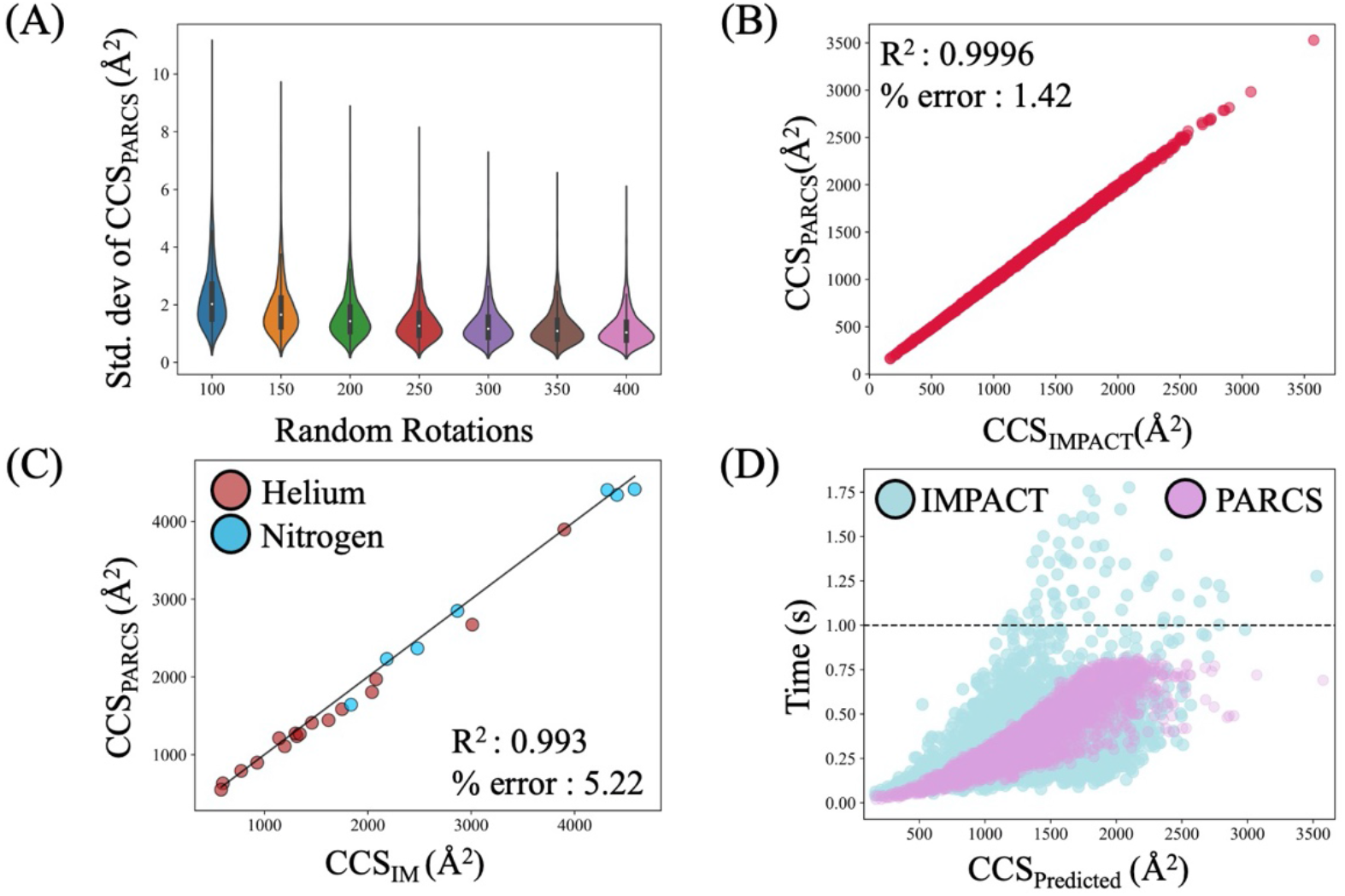
Analysis of PARCS algorithm. (A) Convergence of PARCS calculation over 100-400 random rotations. For more than 250 rotations the mean standard deviation was below 1.5 Å^2^ (B) Comparison of CCS_PARCS_ to that of CCS_IMPACT_ exhibited a very strong correlation (R^2^ of 0.9996). (C) A strong correlation (R^2^ of 0.993) was observed for predicted CCS_PARCS_ values of PARCS when compared with CCS_IM_ from nitrogen (blue) and helium (red) buffer gas for the experimental dataset. (D) Comparison of CCS calculation time of PARCS and IMPACT showed that PARCS and IMPACT performed equally well in terms of speed.

For all proteins in the PARCS evaluation dataset, CCS calculated by PARCS was compared to CCS calculated by IMPACT, one of the most widely used CCS calculation methods, as shown in Figure 2B. A strong correlation (R^2^ = 0.9996) was observed between CCS_PARCS_ and CCS_IMPACT_ with a root mean squared error (RMSE) of 20.38 Å^2^ and an average percent error of 1.42%. The results demonstrate that PARCS calculates CCS values as accurately as other projection approximation methods. CCS_PARCS_ were then compared to CCS_IM_ for the experimental dataset. We observed a strong correlation (R^2^_PARCS_ = 0.993) between CCS_PARCS_ and CCS_IM_ values as shown in Figure 2C, where IM data collected in nitrogen and helium buffer gas are shown in blue and red respectively. We observed an average percent error of only 5.22% (similar to that of IMPACT at 5.61%) from CCS_IM_. To use IM data in computational structure prediction methods (where CCS prediction is required on a large number of decoy structures), the speed of CCS calculations should be within about a second. Therefore, calculation times of PARCS were compared to that of IMPACT as presented in Figure 2D. Using the PARCS evaluation dataset, PARCS took an average of 0.40 seconds to calculate the CCS of proteins when 300 random rotations were used compared to 0.32 seconds for IMPACT. Thus, the timing of PARCS was comparable to IMPACT. We note that the slightly longer average time for PARCS was due to additional steps performed by Rosetta when reading in a PDB file (such as checks for correctness and adding missing hydrogens^52^). For all 4465 proteins, calculations for PARCS completed in under 1.0 second as shown in Figure 2D. These results indicate that PARCS in Rosetta offers similar speed and accuracy to established PA algorithms. We hypothesized that the information contained within CCS_IM_ may be sufficient to predict structures using CCS_PARCS_.

### PARCS in IM score function can improve model selection in ideal dataset (predicted CCS of known structures)

In this study we sought to investigate the usefulness of the structural information encoded in IM data for predicting single-subunit proteins. However, it was unclear whether a single CCS value, encoding overall size and shape, was sufficient to distinguish near-native from incorrect protein models. To test how useful the information in CCS was for structure prediction, an IM score function has been developed to score structures based on the (dis)agreement with experimental IM data. To assess the capabilities of this score function to adequately distinguish good from bad models, we first tested it on the ideal dataset (where the experimental CCS_IM_ was replaced with CCS_PARCS_ of the native structure). For each protein in the ideal dataset 10,000 potential structures were generated using the *ab initio* protocol and scored using the developed IM score function (E_IM_). Prediction results with and without the inclusion of ideal IM data were evaluated and compared based on agreement with experimental structures (using the RMSD of the lowest scoring model, i.e., the predicted structure). We observed a significant improvement in model quality upon the inclusion of ideal IM data. As highlighted in Figure 3A, the RMSD of the predicted structures was improved or unchanged for 82 out of 100 proteins when ideal IM data were included in the scoring. Three of these improvements are shown in Figure 3B, where the native structure (salmon) was compared to the predicted model without and with the inclusion of ideal IM data in blue and pink, respectively.

**Figure 3:**
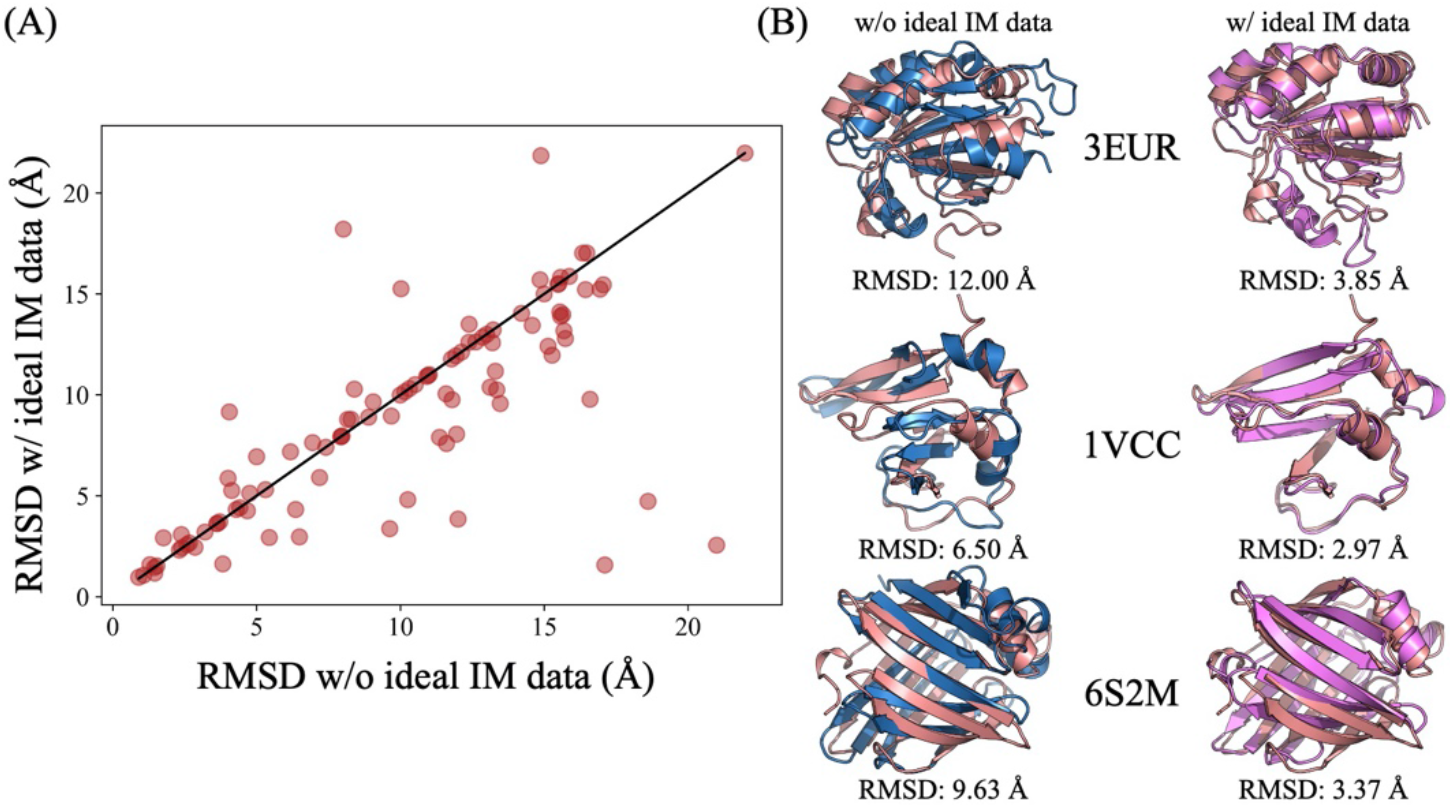
(A) Comparison of the RMSD of the predicted structure of 100 proteins in the ideal dataset. The RMSD improved or remained unchanged for 82 proteins when ideal IM data were used for scoring. (B) Comparison of predicted structures without (blue) and with (pink) the inclusion of ideal IM data to their native structures (salmon).

For the subset of models where the RMSD improved, it improved by an average of 3.62 Å. While the results also suggest that 18 proteins showed an undesirable increase in RMSD (when compared to selection with *E_ROSETTA_*), the average increase in RMSD of this subset was only 2.35 Å. Within these 18 proteins, there were only four cases where the RMSD increased by more than 2 Å. However, three of those four cases were not correctly predicted without the aid of ideal IM data either. The overall average RMSD improved by 0.93 Å for the ideal dataset. Along with the improvement of model selection, this score function also improved definition of the energy funnel, as quantified by the P_near_ metric. From the P_near_ analysis we saw a 7.3-fold increase upon the inclusion of ideal IM data. This suggests that inclusion of IM data significantly improved the goodness of the score versus RMSD funnel. These results indicated that the overall size and shape information contained in the IM data indeed had a strong potential to facilitate the discrimination of good from bad models. Given the sparseness of the data (CCS is a single number denoting the average cross-sectional area of the protein) the improvement was quite significant. While an encouraging proof of principle, these results did not account for the uncertainty associated with real experimental IM data. When experimental IM data are used for the structure prediction, additional uncertainty is included (an average percent error of 5.22% between CCS_PARCS_ and CCS_IM_ was observed for the experimental dataset). Therefore, the effectiveness of IM data to improve structure prediction still needed to be tested on a dataset with experimental IM data.

### IM data improved model selection of protein structures in experimental dataset

For proteins in the experimental dataset, 10,000 decoy models were generated with the *ab initio* protocol for proteins with fewer than 155 residues and comparative modeling (CM) for proteins with more than 155 residues. Each of these decoy models was then scored with IM data (Equation 2) and the predicted models without and with data were compared. Again, we saw a notable improvement in model quality upon the inclusion of IM data. In Figure 4A, the RMSDs of the best scoring models without and with IM data are shown. The RMSD of the predicted models for proteins in the experimental dataset either improved or remained unchanged in all 23 cases. The RMSD improved by an average of 2.01 Å when IM data were utilized as restraints. Of these 23 cases, 15 proteins were ultimately predicted with an RMSD of less than 5.50 Å, compared to 10 proteins without IM data. Furthermore, it was observed that for the subset of proteins where the CM and *ab initio* protocols (shown as triangles and circles respectively in Figure 4A) were used for model generation, the average RMSD improved by 1.61 Å and 2.44 Å respectively. Figure 4B shows structures of the predicted models (aligned to the native models in salmon) obtained without and with the inclusion of IM data (in blue and pink, respectively). The largest improvement was observed for the system β-crystallin B2 (PDB ID: 1YTQ), which improved from 17.7 Å to 5.0 Å. The score vs RMSD distributions for several benchmark proteins before (blue) and after (pink) scoring with IM data are shown in Figure 5. In these distributions, the predicted models without and with IM data are marked with a blue and pink star, respectively. We also observed a general improvement in P_near_ upon scoring with IM data with a 4.5-fold average improvement.

**Figure 4:**
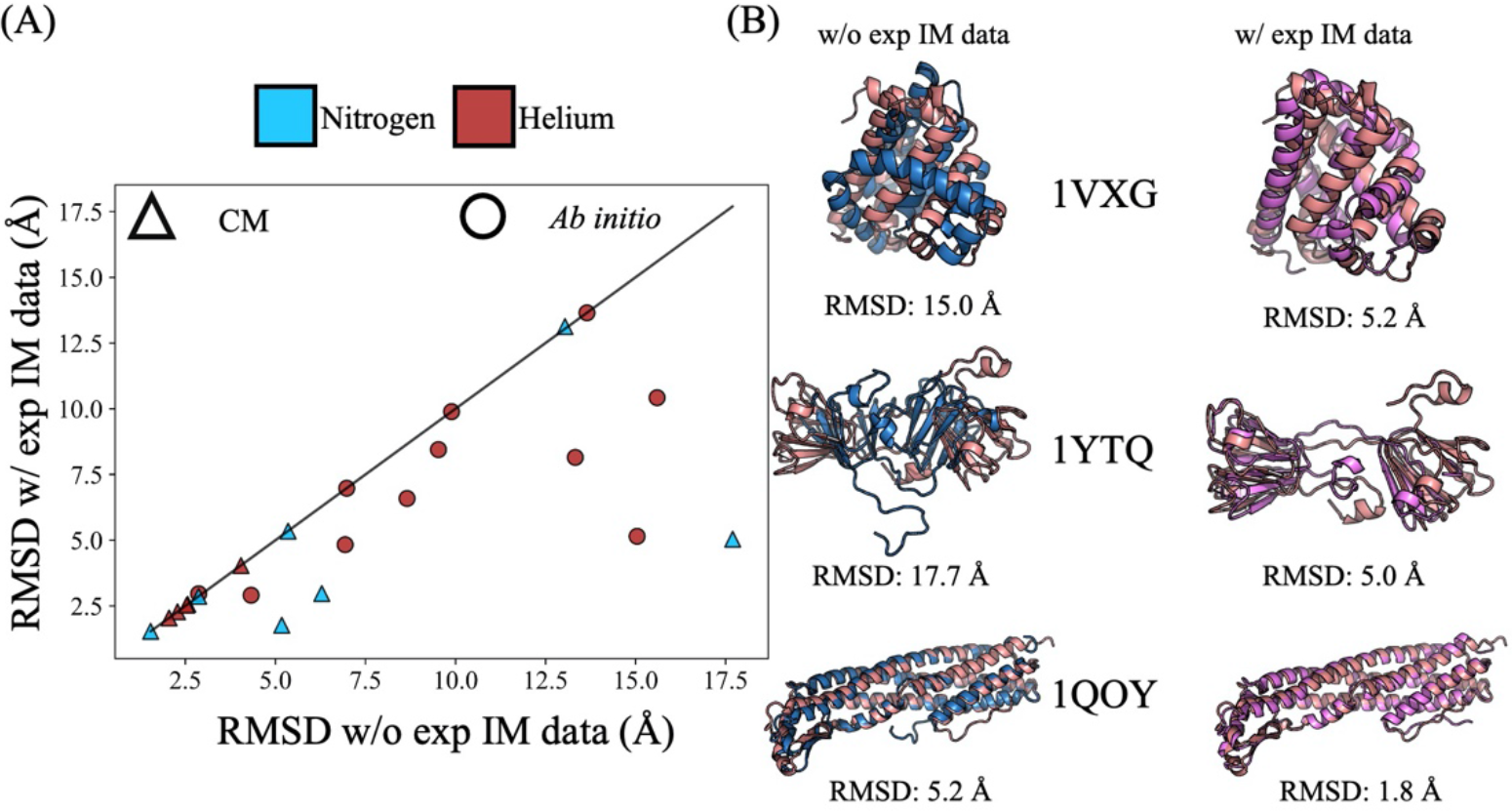
(A) RMSD of the best scoring models with IM data (blue and red for nitrogen and helium buffer gas respectively) when compared to RMSD without IM data for 23 proteins in the experimental dataset. Structures predicted with CM and *ab initio* are represented with a triangle and circle respectively. (B) Comparison of predicted structures without (blue) and with (pink) the inclusion of IM data to their native structures (salmon).

**Figure 5:**
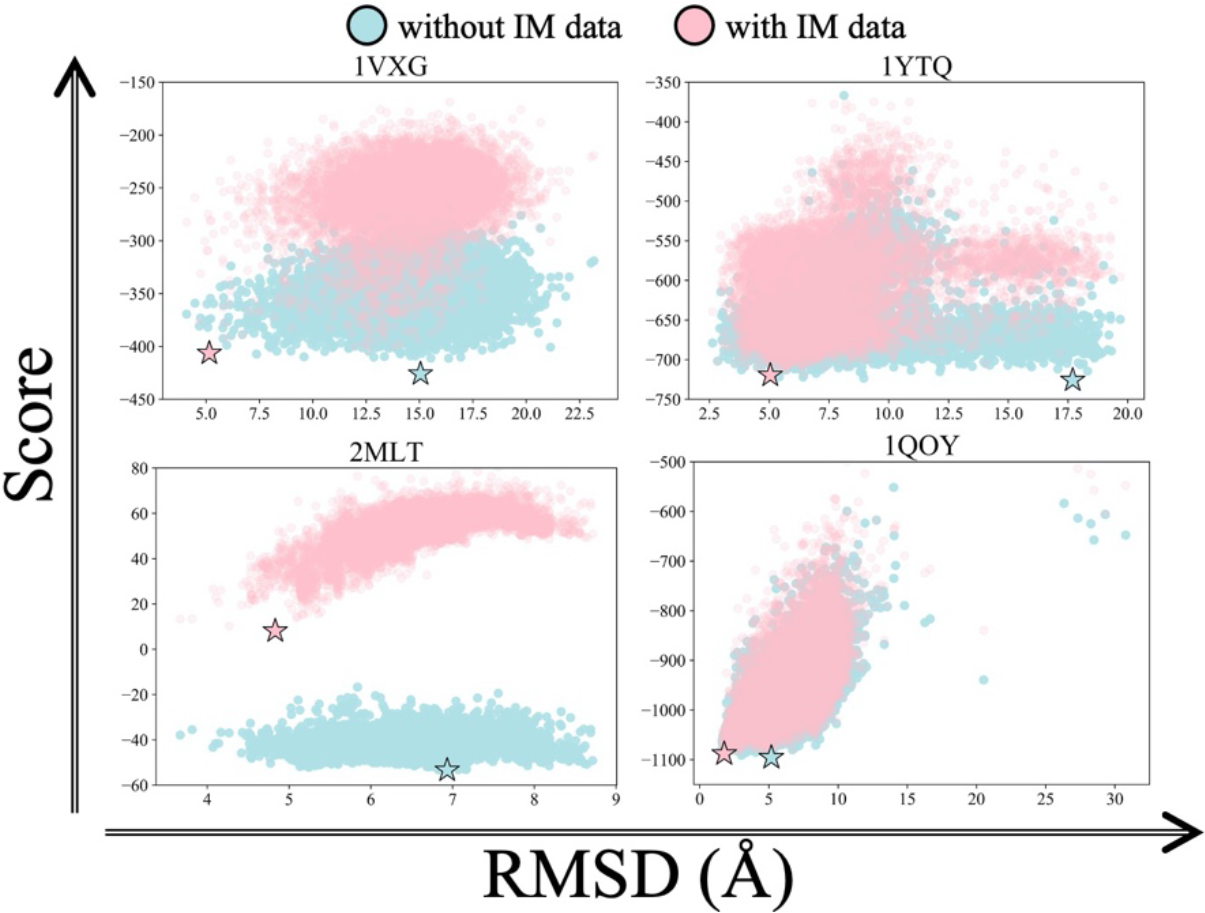
Score vs RMSD plots without (blue) and with (pink) IM data for four proteins that showed significant improvement in the experimental dataset. The best scoring models from both predictions are marked with a blue and pink star, respectively.

Additional analyses on the predicted structures of the experimental dataset further revealed general improvements upon the inclusion of IM data. TM score analysis of this dataset suggested improvements with IM data. We observed, as shown in Table S5, that 74% of the proteins had a TM-score greater than (or equal to) 0.5 when IM data were incorporated as opposed to 65% without the use of IM data. The TM-scores of the best scoring models of the CM subset were generally higher than those of the *ab initio* subset, while the biggest improvements were observed for the *ab initio* subset. General improvements were also seen in a GDT_TS analysis of the best models predicted upon the inclusion of IM data with an average percent score improvement of 2.88%. Similarly, the average GDT_TS of the CM protein subset was higher than that of the *ab initio* subset. In all cases, improvements were observed when IM data were used, with average improvements of 1.51% and 4.39% for the CM and *ab initio* subsets, respectively. All results, as summarized in Table 2, demonstrated that experimental IM data can offer useful information regarding the structural shape of the protein which consequently aids in improved scoring and model selection.

**Table 2:**
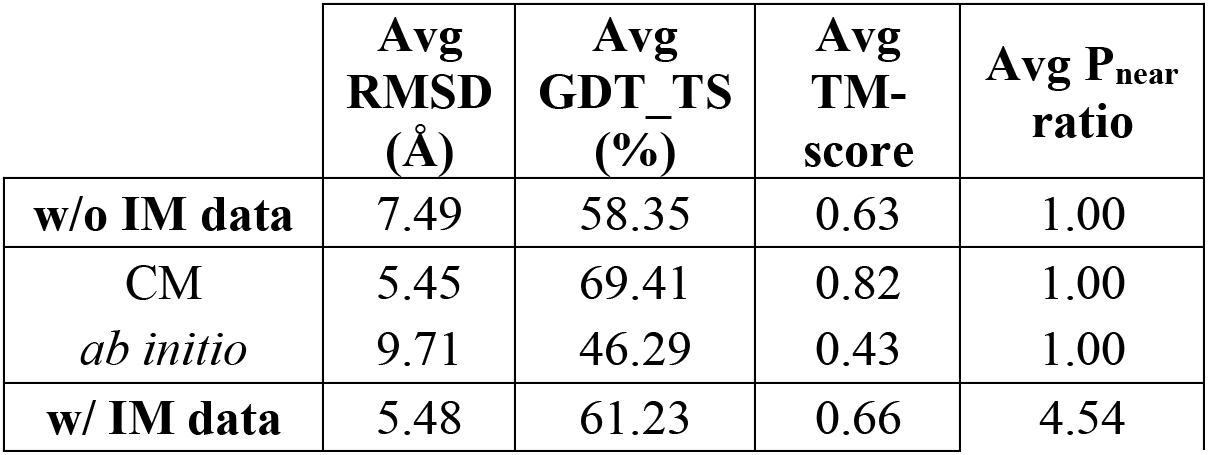
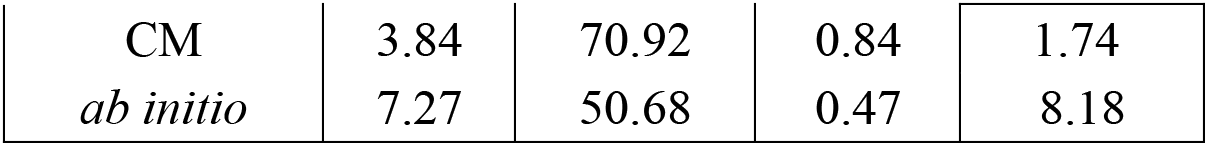
Metric analysis for the best scoring model with and without IM data for the experimental dataset.

| | Avg<br>RMSD<br>(Å) | Avg<br>GDT_TS<br>(%) | Avg<br>TM-score | Avg $P_{\text{near}}$<br>ratio |
| --- | --- | --- | --- | --- |
| **w/o IM data** | 7.49 | 58.35 | 0.63 | 1.00 |
| CM | 5.45 | 69.41 | 0.82 | 1.00 |
| *ab initio* | 9.71 | 46.29 | 0.43 | 1.00 |
| **w/ IM data** | 5.48 | 61.23 | 0.66 | 4.54 |
| CM | 3.84 | 70.92 | 0.84 | 1.74 |
| *ab initio* | 7.27 | 50.68 | 0.47 | 8.18 |

### Confidence measure successfully discriminated accurate and inaccurate models

The inclusion of IM data helped improve structure prediction for all 23 proteins in the experimental dataset. However, there were 8 cases where the RMSD of the selected model (even after improvement) was greater than 5.5 Å. This knowledge was available to us since the native structures were known for the models generated within this benchmark dataset. However, in true blind structure prediction protocols, RMSD information is not available. For this reason, we developed a confidence measure that allowed us to selectively flag successful prediction cases in the absence of native structure. The confidence measure was defined as the average residue score of the top 100 scoring models. According to this metric analysis, the high and low confidence structures were separated by a score cutoff of −2.54. This metric successfully flagged all inaccurate predictions as low confidence, whereas all high confidence predictions were accurate as shown in Figure 6. We saw similar trends when we used this confidence metric to evaluate top scoring models from the ideal dataset as shown in Figure S2.

**Figure 6:**
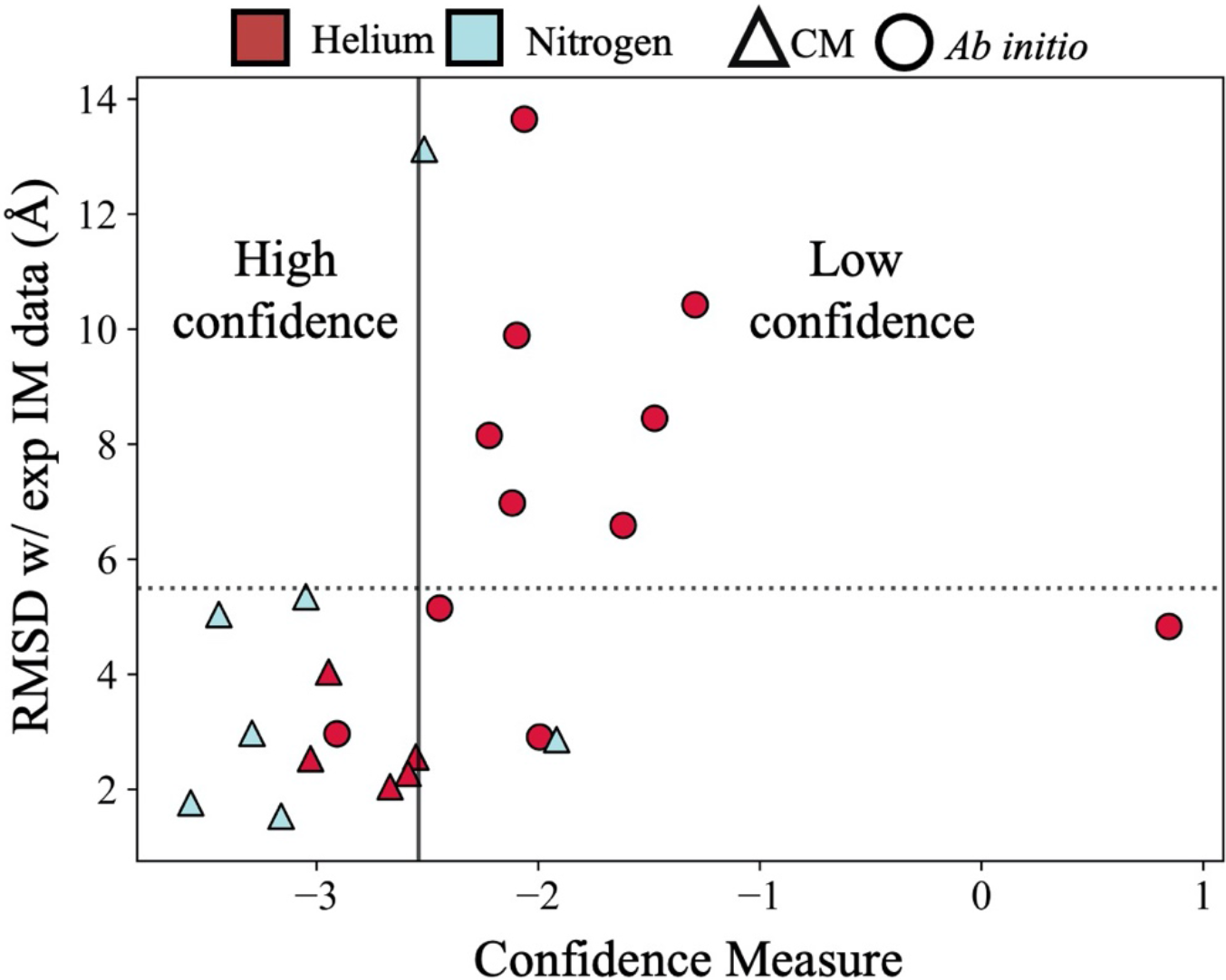
Confidence measure (defined as average residue score of top 100 predicted models) successfully separated the best scoring models from the experimental dataset into two groups, high confidence, and low confidence.

## Conclusion

Ion mobility has emerged as a prime tool to study proteins in their native states using MS. However, the information obtained is sparse, not directly allowing for full structure elucidation. Thus, computational techniques are needed to deduce structural information from IM data. In this study we developed the first such algorithm for structure prediction of single subunit proteins from IM data. To achieve this, we first developed a method (PARCS) that could predict CCS from structures, which has been implemented in Rosetta as a stand-alone application. Following the successful benchmarking of this application, a score term, based on restraints derived from IM data, has been developed to predict native-like structures. This score term was tested on a set of 100 structures from the PDB, where CCS_PARCS_ of the native structure was treated as the experimental CCS_IM_. This was done, as a proof of principle, to check if the score function could translate the structural information (encoded in IM data) into spatial restraints in the absence of model error. Based on RMSD analysis, we observed that the inclusion of IM data improved structure prediction results for 82 out of 100 structures. Following this positive validation, the score function was tested on a benchmark set of 23 proteins with experimental IM data. We showed that IM data improved model selection, as demonstrated by analyzing the best scoring models with several metrics. We also developed a confidence metric to successfully separate good predictions from bad predictions in the absence of native structure. Our current computational workflow illustrates that CCS obtained from IM experiments, despite its sparseness, provides sufficient information on the overall shape and size of proteins to be used as restraints to improve model selection in protein structure prediction. This study also further demonstrates the close connection between the solution and gas phase structure in native IM techniques. Our developed CCS calculation method and score function are freely and easily accessible through Rosetta Commons. A tutorial with examples on how to use the PARCS application as well as the use of IM data in structure prediction has been included in the SI. Further work will focus on improving methods to incorporate CCS data for protein complexes using RosettaDock^74^ and on the use of multiple complementary MS data (such as the combination of CL and/or SID data with IM data) for protein and complex structure prediction in Rosetta.

## Supporting information

Supplemental information for the manuscript and will be used for the link to the file on preprint site

## Acknowledgments

We thank the members of the Lindert lab for many useful discussions. We would like to thank the Ohio Supercomputer Center for valuable computational resources^75^. We also thank Alyssa Stiving and Benjamin Jones for the data they collected on beta lactoglobulin, carbonic anhydrase, ubiquitin and serum albumin. This work was supported by NIH (P41 GM128577, R21A125804). Additionally, integrative protein modeling was supported by a Sloan Research Fellowship to S.L.

---

TOC Figure:

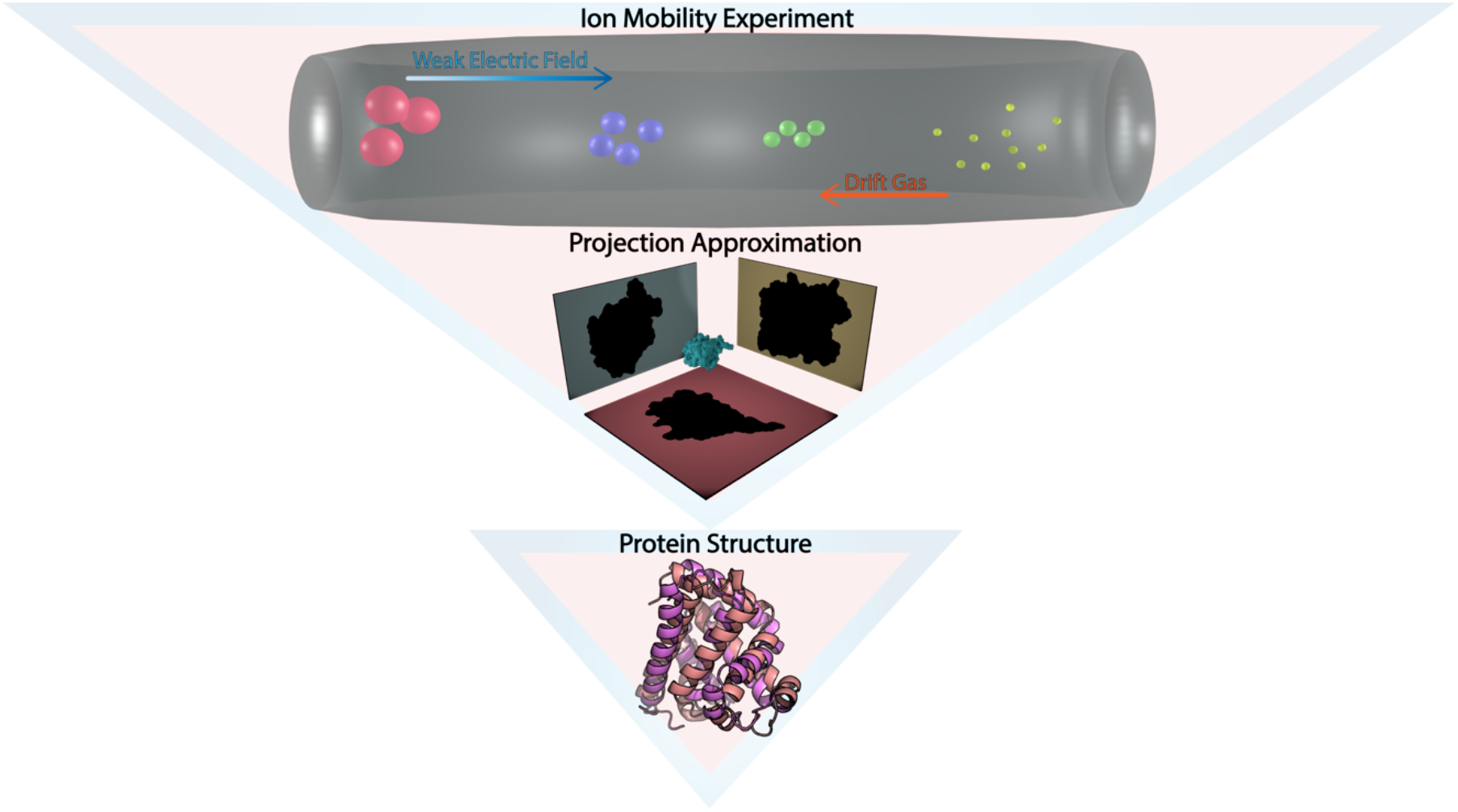

