## Supplemental information for the manuscript and will be used for the link to the file on preprint site for "Protein shape sampled by ion mobility mass spectrometry consistently improves protein structure prediction"

### Supplementary Information

#### Procedures for *ab initio* and comparative modelling protocol in Rosetta

The PDB structures of Lethal Factor N-Terminus (1J7N), Cytolysin A (1QOY), β-crystallin B2 (1YTQ), Fragaceatoxin C (3VWI) and Protective Antigen 63 (4H2A) from the experimental dataset had missing and/or extra residues compared to the protein under experimental IM conditions. For these structures, to ensure that our native structure corresponded to the exact same sequence that was used in the IM measurements, coordinates for missing residues at the C and N termini (CT and NT respectively) were built, as outlined in Table S3, using a modified CM protocol in Rosetta[1-7]. For the ideal and experimental data sets as shown in Table S1 and Table S2 respectively, a decoy set of 10,000 structures was generated for each protein using the AbinitioRelax and comparative modeling (CM) algorithms (as appropriate) within Rosetta. Extensive details about ab initio and CM protocol can be found elsewhere[8]. All fragments were generated using the fragment picker tool[9]. For systems where the ab initio protocol was used (Table S1 and Table S2) for generating the decoy sets, the required fragments were generated by excluding homologs. However, for cases of poor sampling, where the minimum RMSD of the decoy set was greater than to 7 Å, the fragments were re-generated by including homologs. And the decoy set of 10,000 structures was built again with the new fragments. For systems that required the CM protocol (Table S2), templates (and their weights) were chosen (Table S4) such that a broad RMSD distribution was obtained with respect to the native protein. This was done because presence of both native-like and non-native-like models was necessary for benchmarking purposes to demonstrate the ability of CCS data to distinguish between good and bad models. All generated structures were subjected to the Rosetta Relax protocol using the Rosetta energy function (REF2015)[10] and the lowest scoring models were then designated as the final models in all above cases.

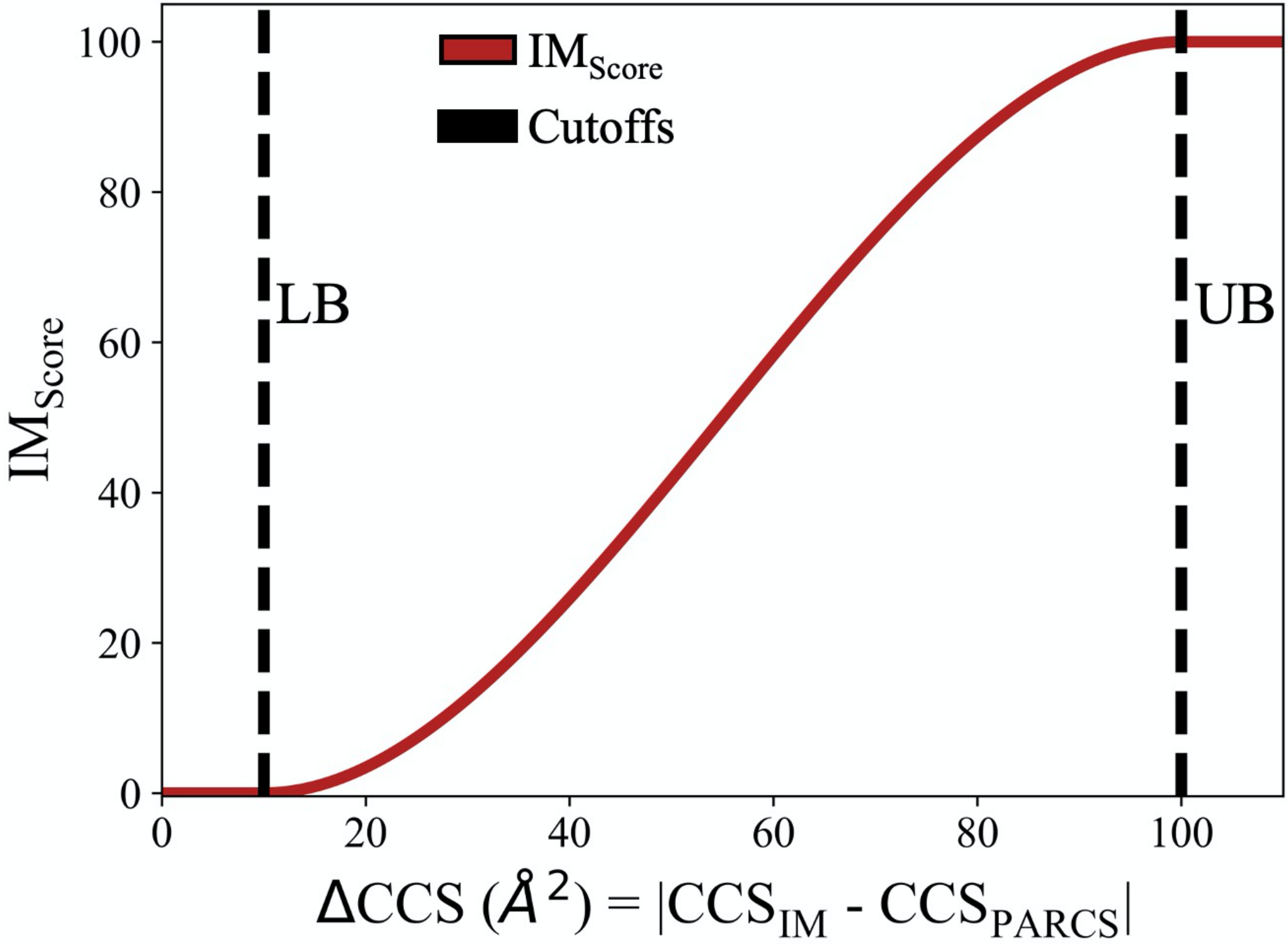


**Figure S1:** *$IM_{Score}$* term is a fade function where LB and UB are the lower and upper bound cutoffs set at 10 $Å^2$ and 100 $Å^2$, respectively. This term penalizes structures based on the absolute difference between the experimental CCS and the structures' predicted CCS.

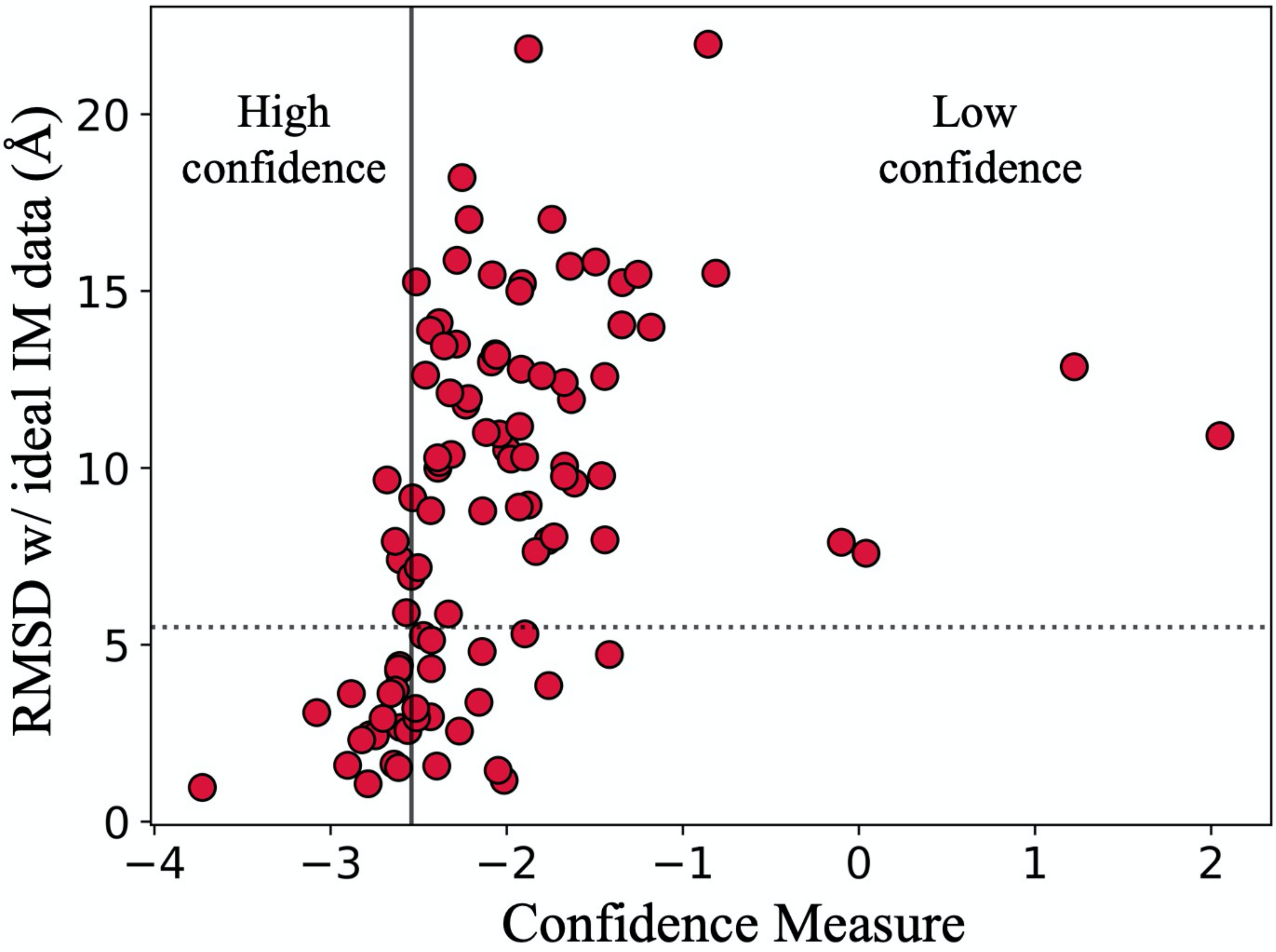


**Figure S2**: Confidence measure (defined as average residue score of top 100 predicted models) successfully separated the best scoring models for the ideal dataset into two groups, high confidence, and low confidence.

**Table S1:** Protein structures from ideal dataset (PDB ID) with ideal $CCS_{IM}$ data (predicted $CCS_{PARCS}$ of native structure), number of residues (**Length**), along with their RMSDs and $P_{near}$ ratio.

| PDB ID | $CCS_{IM}$ (Å$^2$) | Length | Chain ID | RMSD w/o IM data (Å) | RMSD w/ IM data (Å) | $P_{near}$ w/ IM data : $P_{near}$ w/o IM data |
|---|---|---|---|---|---|---|
| 1D0D | 844 | 60 | A | 7.94 | 7.94 | 1.08 |
| 1T8K | 897 | 77 | A | 1.07 | 1.07 | 1.16 |
| 1UFI | 844 | 48 | A | 11.93 | 11.94 | 4.76 |
| 1X2I | 935 | 68 | A | 2.86 | 2.45 | 2.22 |
| 1ZVA | 1086 | 75 | A | 8.02 | 18.21 | 0.92 |
| 2J5Y | 849 | 61 | A | 2.39 | 2.43 | 0.76 |
| 2WX3 | 830 | 43 | A | 11.36 | 7.90 | 208.74 |
| 2XVC | 785 | 56 | A | 2.31 | 2.31 | 1.04 |
| 3A1Y | 794 | 58 | A | 3.82 | 1.63 | 1.37 |
| 3A4R | 963 | 79 | A | 4.41 | 4.41 | 1.04 |
| 3IGM | 1027 | 56 | A | 15.51 | 15.51 | 35.58 |
| 3P1X | 929 | 70 | A | 1.53 | 1.53 | 1.41 |
| 4GKG | 1076 | 52 | A | 21.00 | 2.56 | 2.48 |
| 4MP0 | 839 | 30 | B | 12.86 | 12.86 | 1.00 |
| 4OU0 | 836 | 66 | A | 12.62 | 12.62 | 1.44 |
| 4XYP | 1306 | 70 | A | 21.98 | 21.98 | 1.00 |
| 4ZDT | 932 | 69 | A | 9.68 | 8.95 | 0.90 |
| 5EQ0 | 867 | 55 | A | 8.16 | 8.79 | 0.45 |
| 5H1X | 1030 | 48 | A | 18.61 | 4.72 | 3.02 |
| 5LB7 | 1108 | 78 | B | 1.29 | 1.59 | 1.05 |
| 5N41 | 754 | 60 | A | 9.05 | 9.66 | 0.46 |
| 5XA5 | 595 | 27 | B | 11.60 | 7.59 | 84.35 |
| 5Z7L | 877 | 43 | C | 17.11 | 1.58 | 1.14 |
| 6FF1 | 855 | 69 | A | 1.76 | 2.92 | 1.12 |
| 6FTO | 969 | 66 | A | 4.76 | 5.13 | 0.75 |
| 6I05 | 923 | 76 | A | 3.20 | 3.20 | 1.18 |
| 6MRR | 918 | 68 | A | 2.38 | 3.08 | 1.02 |
| 6RK1 | 809 | 46 | A | 7.97 | 7.97 | 1.09 |
| 1GD8 | 1217 | 105 | A | 15.55 | 14.11 | 0.07 |

| | | | | | | |
|---|---|---|---|---|---|---|
| 1WS8 | 1129 | 104 | A | 1.45 | 1.45 | 1.02 |
| 2H88 | 1349 | 101 | D | 14.88 | 21.85 | 0.44 |
| 2O9U | 1159 | 95 | X | 13.00 | 13.00 | 1.02 |
| 2WKD | 1070 | 79 | A | 13.22 | 13.22 | 0.93 |
| 3D6W | 1245 | 109 | A | 2.66 | 2.66 | 0.40 |
| 3FN2 | 1430 | 97 | A | 13.35 | 10.25 | 8.31 |
| 3G74 | 1102 | 87 | A | 12.39 | 13.50 | 0.01 |
| 3IC5 | 1229 | 115 | A | 7.19 | 5.91 | 1.69 |
| 3LUQ | 1248 | 112 | A | 3.72 | 3.72 | 1.11 |
| 3LYW | 1075 | 86 | A | 10.96 | 10.96 | 2.56 |
| 3RMQ | 1227 | 111 | A | 9.99 | 9.99 | 0.90 |
| 3VZ9 | 1344 | 103 | B | 6.36 | 4.33 | 2.45 |
| 3WOE | 1258 | 104 | B | 15.68 | 13.18 | 1.33 |
| 4IEJ | 990 | 75 | A | 10.15 | 10.15 | 0.78 |
| 4JTM | 988 | 81 | A | 5.00 | 6.94 | 0.94 |
| 4OD6 | 979 | 82 | A | 4.14 | 5.27 | 1.01 |
| 4PWW | 999 | 85 | A | 3.62 | 3.62 | 1.60 |
| 4WLR | 1169 | 99 | B | 10.25 | 4.81 | 2.44 |
| 4YY2 | 1092 | 99 | A | 0.90 | 0.97 | 1.00 |
| 5CEG | 1216 | 103 | B | 17.05 | 15.46 | 0.34 |
| 5D4N | 1273 | 105 | A | 8.89 | 8.89 | 0.75 |
| 5DHD | 1000 | 98 | A | 15.48 | 15.48 | 1.01 |
| 5DIC | 1243 | 115 | A | 3.63 | 3.63 | 1.17 |
| 5EVF | 1224 | 105 | A | 12.12 | 12.12 | 1.16 |
| 5NHU | 622 | 24 | I | 10.91 | 10.91 | 1.07 |
| 5X1E | 1200 | 111 | D | 14.58 | 13.45 | 0.22 |
| 5XPV | 1078 | 100 | A | 13.20 | 12.59 | 0.80 |
| 6AZ5 | 1182 | 116 | A | 15.64 | 13.98 | 0.08 |
| 6C4Q | 1022 | 85 | A | 4.01 | 5.87 | 0.33 |
| 1RG8 | 1301 | 141 | A | 6.95 | 7.63 | 0.36 |
| 1T2W | 1446 | 150 | A | 13.47 | 9.57 | 3.19 |
| 1V4P | 1472 | 151 | A | 16.48 | 17.03 | 0.00 |
| 2CO3 | 1457 | 134 | A | 16.44 | 15.22 | 11.17 |

| | | | | | | |
|---|---|---|---|---|---|---|
| 2Q78 | 1530 | 135 | A | 8.28 | 8.79 | 1.99 |
| 2XQX | 1366 | 147 | A | 14.21 | 14.04 | 0.47 |
| 2YKZ | 1318 | 126 | A | 4.68 | 4.26 | 1.17 |
| 2Z51 | 1639 | 154 | A | 15.00 | 15.00 | 0.02 |
| 3AK8 | 1531 | 154 | A | 4.29 | 4.32 | 2.09 |
| 3EUR | 1350 | 140 | A | 12.01 | 3.85 | 5.18 |
| 3F7X | 1284 | 133 | A | 14.86 | 15.70 | 0.00 |
| 3FH3 | 1619 | 146 | A | 15.57 | 13.89 | 0.01 |
| 3GZR | 1503 | 143 | A | 11.78 | 11.78 | 0.46 |
| 3HM4 | 1381 | 153 | A | 11.95 | 8.05 | 8.22 |
| 3KYJ | 1242 | 129 | A | 2.57 | 2.57 | 3.32 |
| 3NYM | 1563 | 124 | A | 15.27 | 11.97 | 247.70 |
| 3RRI | 1466 | 132 | A | 11.00 | 11.00 | 4.74 |
| 3V7B | 1341 | 147 | A | 15.57 | 15.82 | 0.05 |
| 3X37 | 1477 | 130 | B | 15.73 | 12.79 | 0.63 |
| 4CHM | 1342 | 125 | A | 10.02 | 15.26 | 0.47 |
| 4IUM | 1225 | 127 | A | 15.14 | 12.41 | 2.31 |
| 4OUS | 1286 | 132 | A | 16.94 | 15.24 | 8.01 |
| 4XPL | 1576 | 141 | A | 7.92 | 7.92 | 0.76 |
| 4XPX | 1288 | 130 | A | 6.18 | 7.18 | 0.87 |
| 5DYQ | 1268 | 131 | A | 16.59 | 9.78 | 0.43 |
| 5II0 | 1257 | 99 | A | 10.31 | 10.31 | 1.79 |
| 6BWR | 1710 | 144 | A | 13.12 | 10.38 | 1.14 |
| 6F1K | 1315 | 119 | A | 13.30 | 11.17 | 0.63 |
| 6NE2 | 1230 | 119 | A | 12.38 | 12.60 | 6.06 |
| 2ZPM | 1009 | 86 | A | 7.41 | 7.41 | 1.51 |
| 1K8H | 1663 | 133 | A | 16.34 | 17.03 | 0.06 |
| 1EZG | 823 | 82 | A | 1.46 | 1.17 | 1.07 |
| 6RRV | 1480 | 127 | A | 15.87 | 15.87 | 1.10 |
| 1LWB | 1225 | 122 | A | 10.51 | 10.51 | 0.85 |
| 4G3O | 782 | 53 | A | 5.44 | 2.93 | 1.51 |
| 6SWI | 1518 | 121 | A | 8.40 | 10.29 | 0.28 |
| 3NIR | 625 | 46 | A | 5.31 | 5.31 | 0.94 |

| | | | | | | |
|---|---|---|---|---|---|---|
| 6S2M | 1333 | 133 | A | 9.63 | 3.37 | 2.05 |
| 5B8D | 1060 | 99 | A | 11.59 | 10.05 | 0.34 |
| 1Y6X | 1123 | 87 | A | 4.05 | 9.16 | 1.03 |
| 1VCC | 982 | 77 | A | 6.49 | 2.97 | 1.77 |
| 5K6D | 909 | 77 | A | 11.80 | 9.76 | 1.28 |

**Table S2:** Protein structures from experimental dataset (PDB ID) with experimental $CCS_{IM}$ data, lowest charge state (**z**) that was associated with the experimental CCS data, sequence length (Length), probe radius (**PR**) used for PARCS calculation, along with their RMSDs and $P_{near}$ ratio and the modelling protocol (**Protocol**) that was utilized to generate decoys per structure.

| **Protein Name** | **PDB ID** | **$CCS_{IM}$ (Å$^2$)** | **z** | **Length** | **RMSD w/o IM data (Å)** | **RMSD w/ IM data (Å)** | **$P_{near}$ w/ IM data : $P_{near}$ w/o IM data** | **Protocol** | **PR (Å)** |
|---|---|---|---|---|---|---|---|---|---|
| β Lactoglobulin | 1BEB | 1620 | 6 | 156 | 2.56 | 2.56 | 1.00 | CM | 1.00 |
| Carbonic Anhydrase | 1BN1 | 2080 | 6 | 259 | 2.28 | 2.28 | 1.00 | CM | 1.00 |
| Apo Calmodulin | 1CFD | 1750 | 7 | 148 | 13.33 | 8.15 | 3.41 | *Ab Initio* | 1.00 |
| Chicken Lysozyme | 1DPX | 1313 | 5 | 128 | 9.89 | 9.89 | 0.69 | *Ab Initio* | 1.00 |
| α Chymotrypsinogen A | 1EX3 | 2040 | 7 | 245 | 2.05 | 2.05 | 1.00 | CM | 1.00 |
| Ribonuclease A | 1FS3 | 1300 | 5 | 124 | 6.98 | 6.98 | 0.49 | *Ab Initio* | 1.00 |
| β Densin-2 | 1FD3 | 598 | 3 | 41 | 9.52 | 8.45 | 0.46 | *Ab Initio* | 1.00 |
| Alpha-Lactalbumin | 1HFX | 1342 | 6 | 123 | 4.32 | 2.91 | 2.07 | *Ab Initio* | 1.00 |
| Equine Cytochrome C | 1HRC | 1196 | 5 | 104 | 13.65 | 13.65 | 0.18 | *Ab Initio* | 1.00 |
| β-2-microglobulin | 1LDS | 1142 | 6 | 97 | 15.60 | 10.42 | 0.84 | *Ab Initio* | 1.00 |
| Bovine Ubiquitin | 1UBQ | 930 | 3 | 76 | 2.87 | 2.97 | 1.01 | *Ab Initio* | 1.00 |
| Apo Myoglobin | 1VXG | 1459 | 4 | 153 | 15.04 | 5.15 | 67.50 | *Ab Initio* | 1.00 |
| Meletin | 2MLT | 588 | 3 | 26 | 6.93 | 4.83 | 5.18 | *Ab Initio* | 1.00 |
| Pancreatic Trypsin Inhibitor | 6PTI | 775 | 4 | 56 | 8.65 | 6.59 | 1.80 | *Ab Initio* | 1.00 |
| Bovine Serum Albumin | 4F5S | 3900 | 10 | 583 | 2.53 | 2.53 | 1.01 | CM | 1.00 |
| Ovalalbumin | 1OVA | 3010 | 11 | 386 | 4.04 | 4.04 | 1.00 | CM | 1.00 |
| Lethal Factor N-Terminus | 1J7N | 2480 | 7 | 293 | 13.04 | 13.13 | 0.57 | CM | 1.81 |
| Cytolysin A | 1QOY | 2866 | 10 | 300 | 5.17 | 1.77 | 1.52 | CM | 1.81 |
| β-crystallin B2 | 1YTQ | 2184 | 8 | 204 | 17.70 | 5.04 | 2.47 | CM | 1.81 |
| Fragaceatoxin C | 3VWI | 1838 | 7 | 179 | 2.87 | 2.87 | 1.00 | CM | 1.81 |
| Protective antigen 63 | 4H2A | 4316 | 15 | 569 | 6.29 | 2.97 | 8.32 | CM | 1.81 |

| | | | | | | | | | |
|---|---|---|---|---|---|---|---|---|---|
| HoloTransferrin | 3QYT | 4410 | 14 | 679 | 5.35 | 5.35 | 1.00 | CM | 1.81 |
| Lactoferrin | 1LFG | 4580 | 15 | 691 | 1.53 | 1.53 | 1.00 | CM | 1.81 |

**Table S3:** PDB ID for proteins in experimental dataset. Residues that were added and/or removed from C and N terminal (CT and NT respectively) and the residues are added/removed sequentially from left to right for each terminal.

| Protein | PDB ID | Residues added (CT) | Residues removed (CT) | Residues added (NT) | Residues removed (NT) |
|---|---|---|---|---|---|
| Fragaceatoxin C | 3VWI | SA | - | - | - |
| Cytolysin A | 1QOY | EVPEV | - | T | GILDSMA |
| Protective antigen 63 | 4H2A | G | - | - | E11 - R167 |
| Lethal Factor N-Terminus | 1J7N | - | M264 - S776 | GSHMAGGHGDVGMH VKEKEKNKDENKRKDE | - |
| β-crystallin B2 | 1YTQ | HQRGAFHPSN | - | ASDHQTQAGKPQS | - |

**Table S4:** PDB ID and Chain ID of templates along with their coverage and identity to the PDB ID as well as the weights used for CM protocol in Rosetta for proteins in experimental dataset > 155 residues.

| Protein Name | PDB ID | Templates | Coverage (%) | Identity (%) | Chain ID | Weights |
|---|---|---|---|---|---|---|
| β Lactoglobulin | 1BEB | 1AQB | 62 | 23 | A | 1.5000 |
| | | 2HZQ | 80 | 14 | A | 0.1000 |
| | | 2R73 | 89 | 20 | A | 0.0100 |
| | | 3SAO | 90 | 22 | A | 0.0010 |
| | | 4ROB | 98 | 43 | A | 0.0001 |
| Carbonic Anhydrase | 1BN1 | 5DOT | 26 | 20 | A | 2.0000 |
| | | 1MWA | 23 | 27 | H | 1.0000 |
| | | 3B1B | 86 | 28 | A | 0.1000 |
| | | 5USH | 84 | 37 | A | 0.0100 |
| α Chymotrypsinogen A | 1EX3 | 1KDQ | 53 | 77 | A | 1.0000 |
| | | 3UIR | 100 | 38 | A | 0.1000 |
| Bovine Serum Albumin | 4F5S | 1MA9 | 75 | 23 | A | 2.0000 |
| | | 5OKL | 95 | 35 | A | 0.0010 |
| | | 4ZBQ | 100 | 74 | A | 0.0001 |

| | | | | | | |
|---|---|---|---|---|---|---|
| Ovalalbumin | 1OVA | 1HLE | 86 | 37 | A | 1.0000 |
| | | 2ARQ | 99 | 38 | A | 4.0000 |
| Lethal Factor N-Terminus | 1J7N | 6PSN | 98 | 100 | L | 1.0000 |
| | | 1XFV | 94 | 35 | A | 1.0000 |
| Cytolysin A | 1QOY | 6OGY | 46 | 24 | A | 1.0000 |
| | | 2AP3 | 40 | 26 | A | 1.0000 |
| | | 4PHQ | 98 | 99 | A | 0.1000 |
| β-crystallin B2 | 1YTQ | 1BLB | 100 | 97 | A | 2.0000 |
| | | 1DSL | 75 | 41 | A | 1.0000 |
| Fragaceatoxin C | 3VWI | 2W07 | 10 | 44 | B | 3.0000 |
| | | 1FUI | 44 | 20 | A | 2.0000 |
| | | 5Z06 | 6 | 20 | A | 2.0000 |
| | | 6LCF | 38 | 31 | A | 1.5000 |
| Protective antigen 63 | 4H2A | 2J42 | 73 | 38 | A | 1.0000 |
| | | 3TEY | 100 | 96 | A | 0.1000 |
| HoloTransferrin | 3QYT | 1OVB | 45 | 56 | A | 1.5000 |
| | | 1JNF | 99 | 78 | A | 0.5000 |
| Lactoferrin | 1LFG | 1GV8 | 46 | 57 | A | 2.0000 |
| | | 1B1X | 99 | 75 | A | 0.9000 |

**Table S5**: Metric analysis included template modeling score (TM-score) and global distance test total score (GDT_TS) at a distance cutoff of 5 Å of best scoring structure predicted with and without IM data for proteins in experimental dataset.

| PDB ID | TM-score w/ IM data | TM-score w/o IM data | GDT_TS w/ IM data | GDT_TS wo/ IM data |
|---|---|---|---|---|
| 1BEB | 0.87 | 0.87 | 81.41 | 81.41 |
| 1BN1 | 0.52 | 0.52 | 33.88 | 33.88 |
| 1CFD | 0.40 | 0.38 | 35.81 | 34.80 |
| 1DPX | 0.52 | 0.52 | 49.81 | 49.81 |
| 1EX3 | 0.93 | 0.93 | 84.29 | 84.29 |
| 1FD3 | 0.31 | 0.22 | 43.90 | 34.76 |
| 1FS3 | 0.60 | 0.60 | 56.25 | 56.25 |
| 1HFX | 0.82 | 0.83 | 79.06 | 80.89 |

| | | | | |
|---|---|---|---|---|
| 1HRC | 0.21 | 0.21 | 22.62 | 22.62 |
| 1J7N | 0.72 | 0.74 | 58.80 | 62.92 |
| 1LDS | 0.30 | 0.25 | 29.00 | 26.75 |
| 1LFG | 0.97 | 0.97 | 84.12 | 84.12 |
| 1OVA | 0.88 | 0.88 | 68.64 | 68.64 |
| 1QOY | 0.95 | 0.92 | 89.49 | 85.35 |
| 1UBQ | 0.70 | 0.74 | 75.00 | 79.93 |
| 1VXG | 0.57 | 0.30 | 57.52 | 25.33 |
| 1YTQ | 0.70 | 0.46 | 65.20 | 45.10 |
| 2MLT | 0.44 | 0.44 | 65.39 | 60.58 |
| 3QYT | 0.85 | 0.85 | 66.90 | 66.90 |
| 3VWI | 0.81 | 0.81 | 65.22 | 65.22 |
| 4F5S | 0.93 | 0.93 | 70.03 | 70.03 |
| 4H2A | 0.94 | 0.94 | 83.01 | 85.04 |
| 6PTI | 0.33 | 0.28 | 43.10 | 37.50 |

**Tutorial 1:**
**General usage of PARCS application**

A structure is required to run this application. To use PARCS to predict the CCS of given structure(s), users need to specify the full path to the executable of the PARCS application (<path/to/Rosetta>/main/source/bin/parcs_ccs_application.default.<os><compiler>release). User also need to provide the full path to Rosetta database with the flag *-database*. Next, the structure of the protein for which CCS is to be predicted is specified with *-in:file:s* (or *-in:file:l* for list of structures) in a format readable by Rosetta. Users may choose to specify the number of rotations with *-ccs_nrots* (default is set to 300). Probe radius is set to 1.0 Å by default to predict CCS in helium buffer gas. The other option is to set it to 1.81 Å by using option *-ccs_prad* to predict CCS in nitrogen buffer gas. By default, the application will save the output containing two pieces of information (the name of the structure file and CCS value in Å$^2$) to a file named 'CCS_default.out'. However, users can define the output file name with the flag *-out:file:o*. General usage of the command-line option to run CCS calculation on a single structure is shown below, where variables that need to be specified by users are shown in brackets (< >) and are defined below:

<path/to/Rosetta>/main/source/bin/parcs_ccs_calc.default.<os><compiler>release *-database* <path/to/Rosetta>/main/database *-in:file:s* <structure> *-ccs_nrots* <number_of_rotations> *-ccs_prad* <probe_radius_in_angstroms> *-out:file:o* <output_file_name>

- path/to/Rosetta – Users' path to Rosetta
- os – Name of operating system (linux, mac, etc.)
- compiler – Name of C++ compiler (gcc, clang, etc.)
- structure – Name of structure for which CCS calculation is performed.
- number_of_rotations – Number of random rotations for CCS calculations. Must be an integer. Default is set to 300.

- probe_radius_in_angstroms – Radius of the buffer gas probe. Default is set to 1.0 Å for helium gas. For nitrogen gas please use 1.81 Å.
- output_file_name – User-defined output file name. Default is set to "CCS_default.out".

**Example usage of PARCS application to predict CCS of ubiquitin (1UBQ)**

In this example the CCS of ubiquitin, with a crystal structure available in the PDB (1UBQ), is predicted with PARCS. Note: In this tutorial we will assume that our operating system is Linux and our compiler is gcc.

1. Create a new directory for input and output files and enter this directory

    > mkdir calculate_ccs_for_known_structure && cd calculate_ccs_for_known_structure

2. Download the crystal structure of ubiquitin from https://www.rcsb.org/structure/1UBQ into the calculate_ccs_for_known_structure directory

3. PDB files often contain other useful information, such as water molecules, non-standard amino acids, additional molecules pertaining to experimental conditions, etc. However, this extra information may cause Rosetta to fail if the input structure file is not properly prepared. Fortunately, Rosetta offers a python script (clean_pdb.py) to work around this issue. Use this python script on the PDB file and specify the file name and chain of interest. For ubiquitin this is chain A and the command is.

    > python <path/to/Rosetta>/tools/protein_tools/scripts/clean_pdb.py 1UBQ.pdb A

    Note: The script clean_pdb.py should produce two files. These are 1UBQ_A.fasta and 1UBQ_A.pdb. For this tutorial only 1UBQ_A.pdb file is utilized.

4. Predict the CCS with 250 random rotations and a probe radius of 1.0 and save output file as '1ubq_predicted_ccs.txt' with the following command.

    > <path/to/Rosetta>/main/source/bin/parcs_ccs_calc.default.linuxgccrelease *-database* <path/to/Rosetta>/main/database *-in:file:s* 1UBQ_A.pdb *-ccs_nrots* 250 *-ccs_prad* 1.0 *-out:file:o* 1ubq_predicted_ccs.txt'

5. 1ubq_predicted_ccs.txt contains two pieces of information, the name of the file and CCS value in Å$^2$ as shown below.

| File_Name | CCS_PARCS |
|---|---|
| 1UBQ.pdb | 927.547 |

**<u>Tutorial 2</u>:**

**General usage of scoring (using $E_{IM}$ score function) predicted structures with IM data**

This tutorial explains how to score structures (obtained from *ab initio* or CM protocol in Rosetta) with IM data (using $E_{IM}$ score function). The users need to provide the full path to the score application (<path/to/Rosetta>/main/source/bin/score.default.<os><compiler>release). Full path to database with the flag *-database* is also required. The users also need to specify the structure generated from the prediction protocol with the flag *-in:file:s* (or *-in:file:l* for list of generated structures). The probe radius (required to predict CCS of structures for use in the score function) is set to 1.0 Å by default for helium buffer gas conditions and can be changed to 1.81 Å (with the flag *-ccs_prad*) if the IM experiment was carried out in nitrogen buffer gas conditions. The number of random rotations (*-ccs_nrots*) is set to 300 by default and can be changed as needed. The experimental CCS, derived from IM for the protein of interest, is provided with the required flag *-ccs_exp*. Users also need to specify the patch file (with the option *-score:patch*) *ccs_imms.wts_patch* that calls the $E_{IM}$ score function. A general usage of this score function is shown below, where the variables that need to be specified by users are shown in brackets (< >) and are defined below:

<path/to/Rosetta>/main/source/bin/score.default.<os><compiler>release *-database* <path/to/Rosetta>/main/database *-in:file:s* <structure_from_prediction_protocol> *-ccs_nrots* <number_of_rotations> *-ccs_prad* <probe_radius_in_angstroms> *-ccs_exp* <experimental_ccs_data> *-score:patch* ccs_imms.wts_patch

- path/to/Rosetta – Users' path to Rosetta
- os – Name of operating system (linux, mac, etc.)
- compiler – Name of C++ compiler (gcc, clang, etc.)
- structure_from_prediction_protocol – Structures generated either by using the *ab initio* or CM protocol in Rosetta.
- number_of_rotations – Number of random rotations for CCS calculations. Must be an integer. Default is set to 300.
- probe_radius_in_angstroms – Radius of the buffer gas probe. Default is set to 1.0 Å for helium gas. For nitrogen gas please use 1.81 Å.
- experimental_ccs_data – Experimental CCS value determined from IM experiments (in Å$^2$). If $CCS_{IM}$ is determined from nitrogen buffer gas, set *-ccs_prad* to 1.81 Å. If CCS data is from helium gas, then by default *-ccs_prad* is set to 1.00 Å.

**Example usage of scoring decoy structures (from *ab initio* protocol) of ubiquitin (1UBQ) with IM data**

This tutorial uses ubiquitin (PDB ID: 1UBQ) as an example. A known structure is not required for model generation but providing a native structure will result in RMSD calculation. In this tutorial we will also assume that our operating system is Linux and our compiler is gcc.

1. Create a new directory for input and output files and enter this directory.

    > mkdir score_with_im_data && cd score_with_im_data

2. Download crystal structure of ubiquitin from https://www.rcsb.org/structure/1UBQ in PDB file format.
3. Prepare the file for use with Rosetta.

    > python <path/to/Rosetta>/tools/protein_tools/scripts/clean_pdb.py 1UBQ.pdb A

4. Use Robetta websever (http://old.robetta.org/fragmentsubmit.jsp) to generate the 3mer, 9mer and secondary structure prediction (recommended). Alternatively use fragment picker tool (if set up correctly) to generate these files with this command.

    > <path/to/Rosetta>/tools/fragment_tools/make_fragments.pl -verbose -nohoms 1UBQ_A.fasta

    Note: The fragment picker tool will generate many files, but the files that are utilized from this step are t001_.200.3mers, t001_.200.9mers and t001_.psipred_ss2.

5. Create a flags file (1ubq_abinitio_flags) for the *ab initio* structure generation protocol with these flags.

    *-in:file:fasta* 1UBQ_A.fasta
    *-in:file:frag3* t001_.200.3mers
    *-in:file:frag9* t001_.200.9mers
    *-psipred_ss2* t001_.psipred_ss2
    *-nstruct* 10
    *-abinitio:relax*
    *-use_filters true*
    *-abinitio::increase_cycles* 10
    *-abinitio::rg_reweight* 0.5
    *-abinitio::rsd_wt_helix* 0.5
    *-abinitio::rsd_wt_loop* 0.5

*-ex1*
*-ex2aro*
*-relax::fast*
*-out:file:silent* ./fold_silent.out

6. Run the AbinitioRelax command on the terminal window with this command.
> <path/to/Rosetta>/main/source/bin/AbinitioRelax.linuxgccrelease *-database* <path/to/Rosetta>/main/database @1ubq_abinitio_flags

7. The flag "*-out:file:silent*" in the 1ubq_abinitio_flags file instructs Rosetta to store all 10 structures in a file named fold_silent.out file. When structure generation is complete, run the Rosetta Relax protocol to relax all 10 structures and save them as individual pdbs with this command.
> <path/to/Rosetta>/main/source/bin/relax.linuxgccrelease *-database* <path/to/Rosetta>/main/database *-in:file:silent* fold_silent.out *-in:file:fullatom -relax:quick -nstruct* 1 *-out:prefix* r_

Note: More structures can be generated by increasing the number associated with the flag "*-nstruct*" in the 1ubq_abinitio_flags file.

8. Store the names of all 10 'relaxed' structures (with all output files having the prefix 'r_' because of the flag "*-out:prefix*") generated from Relax protocol with this command.
> ls r_*.pdb > structurelist.txt

9. Run the score application with this command.
> <path/to/Rosetta>/main/source/bin/score.default.linuxgccrelease *-database* <path/to/Rosetta>/main/database *-in:file:l* structurelist.txt *-ccs_nrots* 250 *-ccs_prad* 1.0 *-ccs_exp* 930 *-score:patch* ccs_imms.wts_patch *-in:file:native* 1UBQ_A.pdb

Note: The flag *-in:file:native* is optional and is used for RMSD calculation.

10. The 'default.sc' file produced by the score application contains a lot of information including the IM term (ccs_imms) that contributed to the $E_{IM}$ score (score), RMSD (rms) and the decoy structure (description) that corresponded to this information as shown below.

| SCORE: | score | ... | ccs_imms | ... | rms | description |
|---|---|---|---|---|---|---|
| SCORE: | -182.443 | ... | 7.046 | ... | 5.691 | r_F_00000005_0001_0001 |
| SCORE: | -145.027 | ... | 31.242 | ... | 11.382 | r_F_00000006_0001_0001 |
| SCORE: | -187.302 | ... | 12.786 | ... | 3.204 | r_S_00000001_0001_0001 |
| SCORE: | -188.296 | ... | 11.096 | ... | 4.594 | r_S_00000002_0001_0001 |
| SCORE: | -105.458 | ... | 71.117 | ... | 9.342 | r_S_00000003_0001_0001 |
| SCORE: | -183.859 | ... | 0.018 | ... | 4.850 | r_S_00000004_0001_0001 |
| SCORE: | -206.509 | ... | 0.000 | ... | 4.049 | r_S_00000007_0001_0001 |
| SCORE: | -40.841 | ... | 100.000 | ... | 11.913 | r_S_00000008_0001_0001 |
| SCORE: | -44.949 | ... | 100.000 | ... | 9.460 | r_S_00000009_0001_0001 |
| SCORE: | -137.448 | ... | 15.016 | ... | 3.404 | r_S_00000010_0001_0001 |

Note: Other terms not important to this tutorial are represented as '…'

11. The $E_{IM}$ score and RMSD of each 'relaxed' structure compared to the native of the generated structure is extracted from the 'default.sc' file (produced by score application) with the following command.

```
> python
        > import numpy as np, pandas as pd, matplotlib.pyplot as plt
        > score_file  = pd.read_csv('default.sc', sep='\s+', header=0)
        > score        = score_file['score']
        > rmsd         = score_file['rms']
        > plt.figure()
        > plt.scatter(rmsd,score,color='pink')
        > plt.xlabel(r'RMSD ($\AA$)')
        > plt.ylabel('Score with experimental IM data')
        > plt.savefig('1UBQ_SCORE_VS_RMSD.png',dpi=300)
        > plt.close()
```

12. View results in file '1UBQ_SCORE_VS_RMSD.png'.

Note: This plot is only meaningful when a large number of structures are generated.
